# Lack of redundancy between electrophysiological measures of long-range neuronal communication

**DOI:** 10.1101/2020.07.16.207001

**Authors:** Daniel Strahnen, Sampath K.T. Kapanaiah, Alexei M. Bygrave, Dennis Kätzel

**Affiliations:** Institute of Applied Physiology, Ulm University, Ulm, Germany; Department of Neuroscience, Johns Hopkins University, USA

**Keywords:** schizophrenia, hippocampal-prefrontal coherence, wPLI, phase-amplitude coupling, spike-phase coupling, Granger causality, MRL, amplitude cross-correlation, theta, *Gria1*

## Abstract

Communication between brain areas has been implicated in a wide range of cognitive and emotive functions and is impaired in numerous mental disorders. In rodent models, various functional connectivity metrics have been used to quantify inter-regional neuronal communication. However, in individual studies, typically only very few measures of coupling are reported and, hence, redundancy across such indicators is implicitly assumed. In order to test this assumption, we here comparatively assessed a broad range of directional and non-directional metrics like coherence, weighted Phase-Lag-Index (wPLI), Granger causality (GC), spike-phase coupling (SPC), cross-regional phase-amplitude coupling, amplitude cross-correlations, and others. We applied these analyses to simultaneous field recordings from the prefrontal cortex and the ventral and dorsal hippocampus in the schizophrenia-related *Gria1-*knockout mouse model which displays a robust novelty-induced hyperconnectivity phenotype. We find that across such measures there is a considerable lack of functional redundancy. While coherence and GC yielded similar results, other measures, especially wPLI and SPC, often produced deviating conclusions. Bivariate correlations within animals revealed that virtually none of the metrics consistently co-varied with any of the other measures across the three connections and two genotypes analysed. Parametric GC showed the qualitatively highest degree of redundancy with other metrics and would be most suitable for connectivity analysis. We conclude that analysis of multiple metrics is necessary to characterise functional connectivity.

## Introduction

Communication between different brain regions is vital for cognition and emotion, and is impaired in a variety of neurological and psychiatric disorders, including schizophrenia and depression. In order to better understand interregional communication in health and disease at the electrophysiological level in rodent models, local field potentials (LFPs) and sometimes action potentials (spikes) are typically recorded from two or more brain areas simultaneously in awake subjects. Subsequently, some metric of interdependency of the signals from two regions are computed.

For example, an influential hypothesis known as *communication through coherence* (CTC) states that information exchange between two connected brain areas depends on the timing of the arrival of incoming activity in a specific phase of a certain network oscillation [1–4]. This is because such network oscillations represent the integrated synaptic activity, and thus peaks vs. troughs represent times of low vs. high probability of synaptic inhibition and, hence, excitability by an incoming long-range input. Therefore, coherence measures a synchrony of oscillations in a certain frequency range and with a certain phase-shift that may allow the activity generated in one region to optimally affect the activity of another region.

In general, measures of phase synchronization aim to quantify if two signals have a consistent phase relationship between each other. Despite being widely used, coherence is prone to confound by volume conduction [5, 6]. Therefore, alternative measures of phase synchronization have been suggested. Nolte et al. demonstrated that using only the *imaginary component of coherence* (ImC) effectively reduces the influence of a volume-conducted signal originating from a common source [7]. Alternatively, the *phase lag index* (PLI) may be used, which disregards the magnitude of the phase lag between signals from two brain regions but evaluates if they differ from a symmetrical distribution [8]. The weighted PLI (wPLI), in turn, applies the combined advantages of the ImC and PLI by taking the detected phase lead or lag and weighing it by the magnitude of the ImC [9]. A constraint related to measures of phase synchronization like the ImC, PLI and wPLI is sample-size bias which usually increases with a lower number of samples. Therefore Vinck et al. additionally introduced a debiased estimator of the wPLI which is more independent from sample size and thus has a higher statistical power than previous measures [9]. For the sake of clarity, the debiased wPLI will be referred to as wPLI throughout this study. Like coherence, it ranges from zero (no synchrony) to one (total synchrony).

It should be noted that several other measures for phase synchronization exist, such as the phase-locking value (PLV) [10] and pairwise phase consistency (PPC) [11]. These measure the constancy of the difference between the instantaneous phases of two signals obtained either by applying hilbert or wavelet transformation and quantifying the distribution of phase differences across observations either by taking the vector average or by determining the distribution of phase differences, respectively.

The stated measures of phase synchronization are attempts to quantify *functional* or *non-directed* connectivity. This means that the quantification of coupling is essentially based on correlation analysis, ignores its temporal structure, and assumes no direction of the influence from one region to another [12, 13]. However, LFP data can also be used to measure *effective* or *directed connectivity* [13]. These are parameters that quantify the potentially causal influence that the activity in one region exerts on another region by taking recurring pairwise patterns in the time series obtained from both regions into account. A computationally simple measure to detect directionality between two time series is *cross-correlation*. That means that correlations are calculated as the LFP-signals are incrementally shifted against one another to obtain a cross-correlation as function of temporal shifts (lags). Adhikari et al. developed a method termed *amplitude cross-correlation* or *cross-amplitude coupling* in which the instantaneous amplitudes of two oscillatory signals filtered in a certain frequency range are cross-correlated to determine if one is leading or lagging the other [14]. If the lags at which the peak of the amplitude cross-correlation function occur is significantly different from 0 ms, it is indicated that one region leads the other one with a certain consistency, which could be due to a directional influence from the leading onto the lagging region. This method was able to identify directional connectivity in the brain related to working memory and fear processing [15–17].

A different measure of directed influence is *Granger causality* (GC). It is a measure to infer directional causation based on the notion that one signal is helpful in predicting the other by statistically comparing results of autoregressive models (ARM). Here, two separate ARMs are calculated: A univariate ARM, where the signal is predicted by a weighted combination of its own past values, and a bivariate ARM where the signal is additionally predicted by the second signal. If the inclusion of the bivariate AR leads to a reduction of variance of the predicted signal, one signal is said to Granger-cause the other [18]. The mathematical foundations of GC and its application to neuroscience has been reviewed extensively elsewhere [18–21]. It should be noted that a non-parametric approach to estimate Granger Causality was shown to be equivalent to the parametric approach we use here [22].

Other indicators of inter-regional communication that partly circumvent problems caused by volume conductance, and are typically interpreted as indicating a causal directional influence include those that measure *different types* of neuronal activity in the different regions; i.e. a low-frequency LFP oscillation (usually in the theta-range) in the presumed dominating region and a local and high-frequency activity is at the receiving end. In contrast to the metrics introduced before, historically, such measures were introduced by way of an actual *biological* discovery of such coupling phenomena, rather than by *a priori* mathematical considerations on how to best assess inter-regional communication. One option is to quantify the extent to which oscillations of *distinct* frequencies are coupled to each other, a phenomenon called cross-frequency coupling (CFC, [23]). Particularly, local phase-amplitude coupling (PAC) - the statistical relationship between the phase of a low-frequency and the amplitude of a high-frequency component - plays an important role in memory processing in the hippocampus of rats [24, 25] and humans [26]. However, *cross-regional* PAC between the hippocampus and prefrontal cortex has also been used and was associated with directed information flow and cognitive functions [6, 27–29]. Since high-frequency brain oscillations mainly reflect local aspects of information processing and low-frequency brain rhythms are relevant for inter-regional communication, CFC might represent a mechanism of transferring information from large scale neuronal networks to local processes [23, 30].

Another very widely used measure is based on the recording of spikes in one (potentially the *influenced*) region alongside the LFP in another (potentially the *influencing*) region. Spikes are generally not considered to be confounded by volume conductance or referencing, and they represent a more direct readout of the actual neuronal activity of a region. Phase-locking of neuronal firing to theta-frequency hippocampal oscillations was shown for example in the PFC [31, 32], entorhinal cortex [33] and the amygdala [34]. E.g. action potentials in these brain regions occurred rhythmically at the same phase of the hippocampal theta rhythm. Such spike-phase coupling (SPC) was observed to correlate with performance in multiple cognitive tasks [31, 32, 35], and has been used to evaluate coupling deficits in mouse models related to schizophrenia [36–38].

The above mentioned measures have been widely used for two decades to assess inter-regional neuronal communication in rodents during a variety of cognitive tasks and disease-related manipulations, typically involving recordings from the hippocampus and prefrontal cortex [31, 36, 37, 39], but also increasingly from the thalamus [40], and the amygdala [41].

However, typically only one or two measures of coupling are calculated and interpreted as sufficient surrogate to quantify task- or manipulation-related differences in actual information exchange between the analyzed regions. In this analytical set-up, the contingency of the achieved conclusions on the choice of the coupling measure are usually not evaluated, but the redundancy of the various measures is implicitly assumed. Likewise, the dependence of the conclusions on the exact placements of electrodes within the analyzed regions and the choice of reference are often not evaluated either. While this might not be regarded as a major concern when an instant of coupling is actually discovered, it certainly presents a problem when interpreting negative data, i.e. the supposed absence of coupling.

We therefore sought to evaluate the redundancy and contingencies of such coupling metrics. To this end we recorded data during a simple behavioural assay – novelty-induced locomotion and its habituation over time – in *Gria1*^−/−^ (KO, knockout) mice and their littermate controls. We have recently shown, that the *Gria1*-KO model, which is related to schizophrenia, shows profound and state-dependent aberrations of hippocampal-prefrontal coupling in this task [42]. We focused on the most widely used connectivity measures – coherence, ImC, wPLI, cross-amplitude coupling, parametric GC, SPC and SPC-related directionality – with respect to three ‘litmus tests’ for redundancy: (a) detection of KO-related alterations of coupling across the 10 min test, (b) detection of KO-related changes of a measure over time, and (c) bivariate within-animal correlation of the analyzed measures. We analyse the widely studied connections between the medial prefrontal cortex (PFC) and the hippocampus – both the dorsal (dHC) and the ventral (vHC) partition - and study four common frequency bands, delta (δ, 1-4 Hz), theta (θ, 5-12 Hz), beta (β, 15-30 Hz) and low-gamma (γ, 30-48 Hz).

## Results

### Elevated locomotor activity in *Gria1-KO* mice during measurement of interregional communication

In order to measure inter-regional coupling we implanted 20 adult *Gria1*^−/−^ mice and 12 littermate controls unilaterally with LFP electrodes in 4 regions, PFC (2 electrodes), mediodorsal thalamus (MD, 1 electrode), dHC (1 electrode) and vHC (2 electrodes), and inserted screws for ground and reference above the cerebellum and frontal cortex, respectively (Fig. 1A). Recordings from all sites were made during a 10 min test of novelty-induced locomotor activity which confirmed the strongly elevated behavioural activity and failure of its short-term habituation over time, as observed in *Gria1*^−/−^ mice before (Fig. 1B, C; 42]). After experiments were completed, the placement of electrodes were evaluated through electrolytic lesion sites and misplaced electrodes were excluded from the dataset; data from the MD was disregarded for most of the subsequent analysis because of the low number of animals with accurate placements. In accordance with our previous study in this mouse line [42], we recorded and analysed all data as referenced to the ground screw above the cerebellum by default and used the data from the frontal reference screw for a separate analysis (displayed in Fig. 7). We extracted LFP signals (Fig. 1D) and multi-unit-activity (MUA) spikes (Fig. 1E) from all depth electrodes. LFPs were filtered in the respective frequency bands for all assessed connectivity measures (Fig. 1F-H). For PAC and SPC the theta-phase angle was extracted using a Hilbert-transform (Fig. 1G, H). Additionally, we sorted the LFP power values obtained from each electrode in distinct frequency bands according to the placement of electrodes in different subdivisions of the PFC (PrL, Cg1 and Cg2), dHC (apical dendritic layers of CA1, CA1 pyramidal cells, CA1 stratum oriens) and vHC (apical dendritic layers of CA1/CA3, CA1 pyramidal cells, dentate gyrus). While we did not conduct statistical analysis given the much smaller number of sites outside the target region (PrL in PFC and apical dendritic layers, including fissure, in the hippocampus), a qualitative inspection suggested that the placements inferred from lesion sites did not noticeably alter the obtained spectral LFP power (Fig. 1I-K).

**Figure 1.**
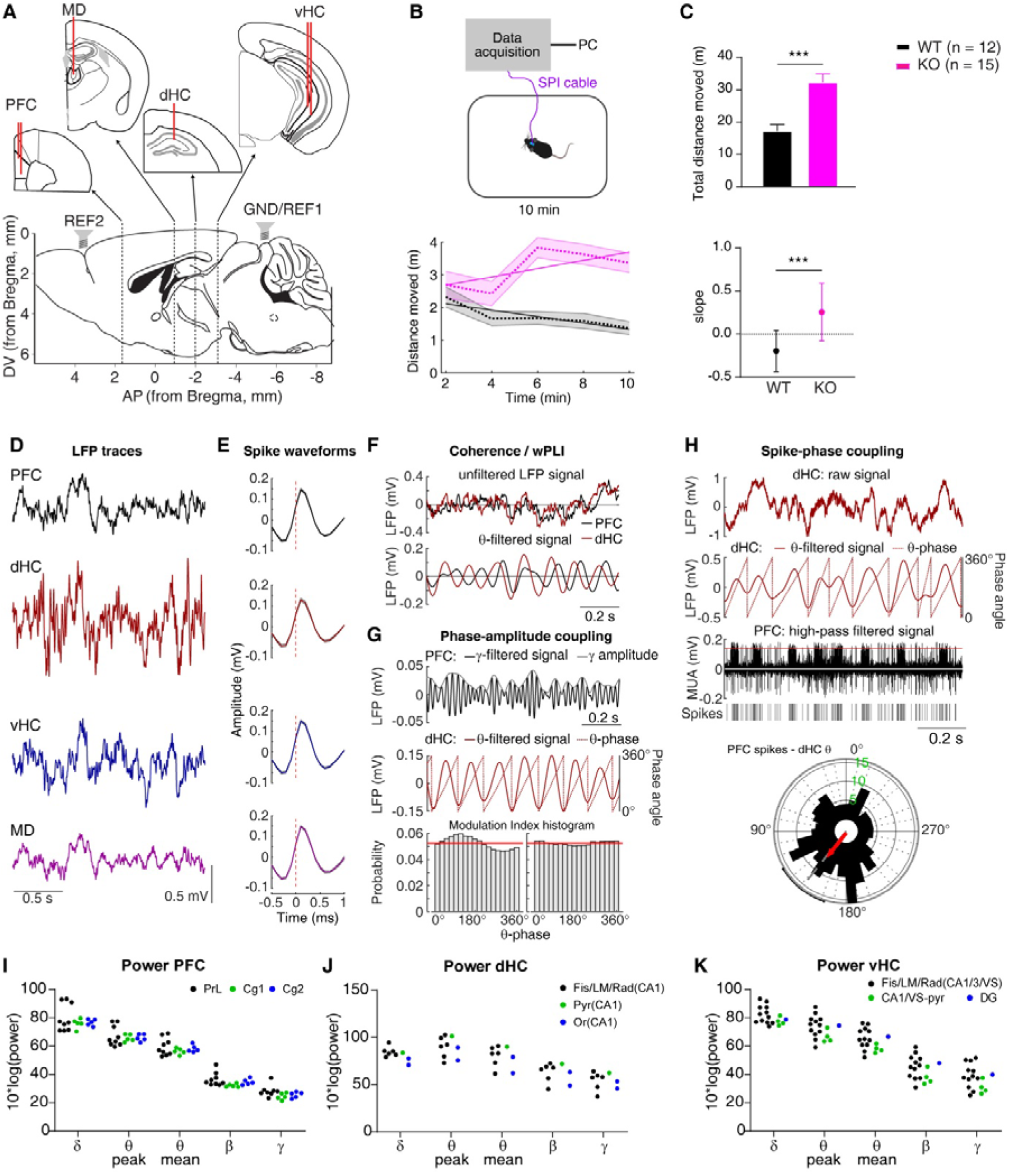
Experimental set-up, behaviour, and recorded signals. (**A**) Placement of LFP and screw electrodes, note that in the majority of mice a bundle of two electrodes was implanted into the PFC and the vHC with an approximate distance of 0.5 mm between electrodes. (**B**) Top, experimental set-up; bottom, distance moved in 2 min bins by *Gria1*^−/−^ (KO, purple) and wildtype controls (WT, black), dashed line indicating mean, shaded region representing SEM, solid overlaid line representing linear interpolation across 2 min bins within each group. (**C**) Same data as in (B) but displayed as total distance moved in 10 min (top) and slope of the interpolated line (bottom). (**D**) Examples of unfiltered LFP traces recorded in the four brain regions. (**E**) Average waveforms of spikes extracted from high-pass filtered signals (MUA) in the regions identified in (D). (**F**) Illustration of the processing for connectivity measures using the same frequency in both regions; raw LFP signal (top) and LFP signal filtered in a specific frequency-range (bottom). (**G**) Illustration of cross-regional θ-γ PAC, whereby the signal in one region is filtered in the low-γ range and the amplitude is extracted (top), while the signal in the other regions is filtered in the θ-range and Hilbert-transformed to extract the θ-phase (middle). The coupling-strength is derived as modulation index (MI) measuring the phase-related change of γ -amplitude (bottom). (**H**) Illustration of MRL-calculation; the raw LFP signal in one region (top) is filtered in the theta-range and Hilbert-transformed to extract the instantaneous phase angle (below, brown); the high-pass-filtered signal in the other region reveals MUA from which spikes are extracted by thresholding (below, black); a circular histogram is computed by assigning each extracted spike to the theta-phase angle at which it occurs and the average of all vectors formed by all spikes in this phase space is calculated as mean resultant vector (red) of which its length (MRL) is taken as indicator of spike-phase coupling strength (bottom). (**I-K**) Power of LFP in the indicated frequency bands (x-axis) and region (top of panel) displayed for each individual electrode that contributed to the WT-dataset colour-coded by the sub-division in which it was placed; hippocampal layers: pyramidal (Pyr), stratum oriens (Or), lacunosum-moleculare (LM), radiatum (Rad), fissure (Fis). No statistical analysis was done given that some placements occurred very rarely.

### Differences in detecting elevated inter-regional theta-range coupling in *Gria1-KO* mice across connectivity measures

We first analysed phase-synchronization along the two prefrontal-hippocampal connections (PFC-dHC and PFC-vHC) and within the hippocampus (vHC-dHC) using coherence and wPLI (Fig. 2A-L). We confirmed our previous observation [42] that PFC-dHC theta coherence is strongly elevated in *Gria1*-knockouts in a novel environment and further increases with time, mirroring the spatial exploration behaviour of this genotype (Fig. 1B, C, Fig. 2A, D). However, this phenotype was by no means specific to the PFC-dHC coupling, but also re-appeared in the PFC-vHC and vHC-dHC connections suggesting a broader deficit of excessive theta-range connectivity. Reassuringly, the same phenotype was revealed by the wPLI metric across connections (Fig. 2G-L). However, when inspecting the other frequency bands, findings were not particularly consistent between the two measures. While both indicators revealed a reduced gamma-range PFC-dHC coupling, a sole analysis with wPLI suggested further differences in the delta (PFC-dHC, vHC-dHC), beta (PFC-dHC), and gamma (PFC-vHC) ranges that would have gone undetected, if using the coherence metric (Fig. 2D-F, J-L). Also, qualitatively, both measures resulted in spectra with quite different shape.

**Figure 2.**
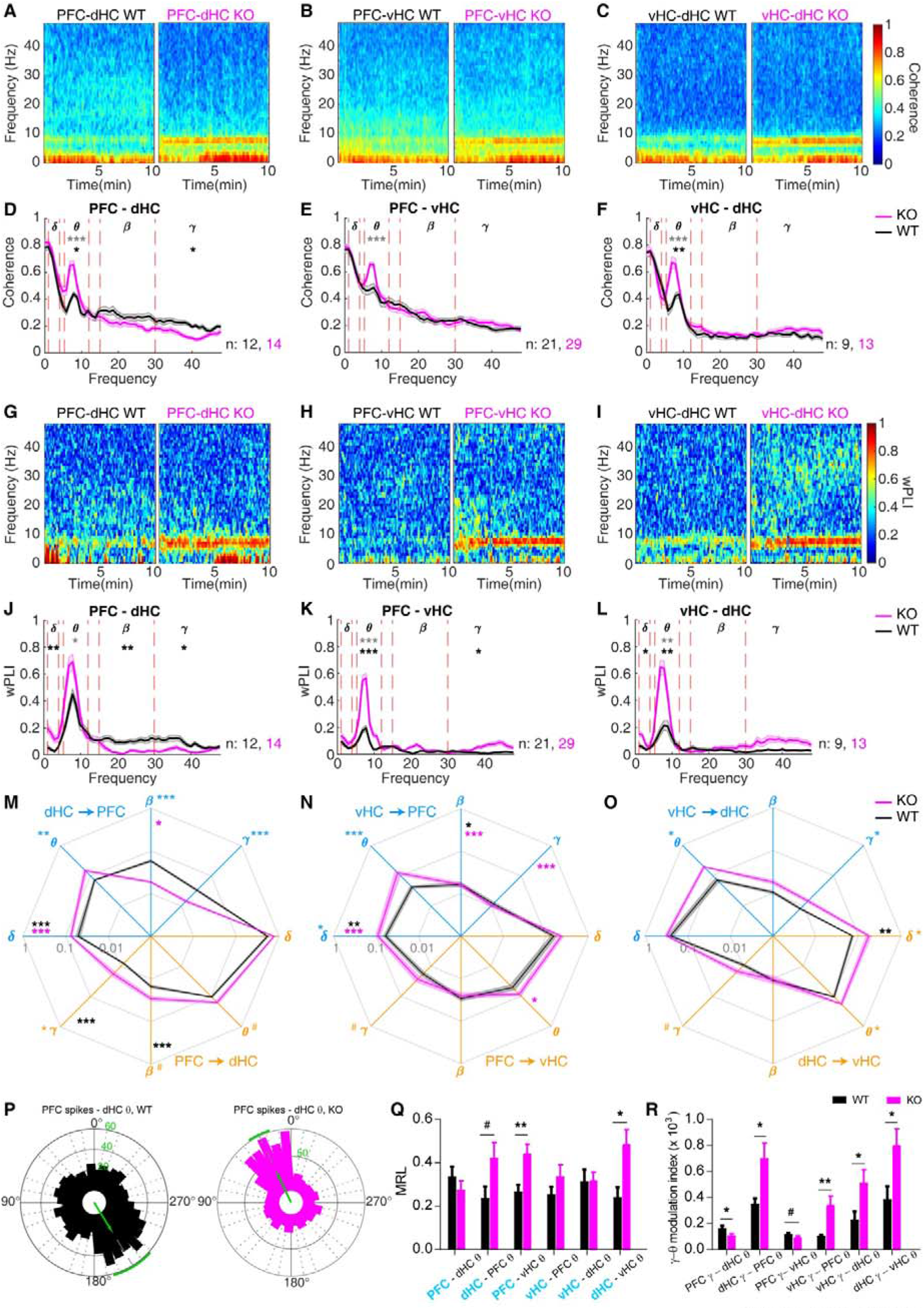
Multi-parameter assessment of inter-regional coupling strength in *Gria1*^−/−^ and wildtype controls across 10 min novelty-induced activity. (**A-L**) Spectrogramms (A-C, G-I) and frequency-spectra (D-F, J-L) displaying coherence (A-F) and wPLI (G-L) along the PFC-dHC (A, D, G, J), PFC-vHC (B, E, H, K) and vHC-dHC (C, F, I, L) connections. Red lines in spectra indicate the boundaries of the analysed frequency bands named by the greek letters at the top. Stars indicate significant differences between genotypes (*t*-test) in mean (black) or peak (grey) coherence (D-F) or wPLI (J-L). (**M-O**) GC in the frequency bands indicated by greek letters and along the directional connections identified by the colour (blue: dHC➔PFC (M), vHC➔PFC (N), vHC➔dHC (O); orange: reverse of the before). Statistical indicators in the same colour identify a difference between genotypes (Sidak post-hoc test, conducted after significant effect of genotype or interaction in a given connection and frequency band in RM-ANOVA); statistical indictors in black (WT) or purple (KO) refer to a significant difference between the GC-values of the two opposing directions within the colour-coded genotype whereby the location of the indicator identifies the direction with *smaller* average GC. Grey numbers refer to GC, plotted as log_10_-scale. All thick lines in D-F and J-O represent the mean value, while shades in lighter opacity represent SEM. (**P**) Example of spike-phase histograms and mean resultant vector (green) for an individual WT (left) and KO (right) mouse illustrating coupling of PFC-spikes to dHC-theta oscillations. (**Q**) Mean resultant vector length (MRL) as indicator of spike-phase locking (as illustrated in P) for 2 directions along the 3 connections, as indicated (blue font identifies the region that contributes the spikes). (**R**) Theta-gamma cross-regional PAC for the named directional connections (first region contributing gamma-oscillations). Significance indicators in (Q-R) refer to group differences (*t*-test); bars display mean ± SEM. # *P* < 0.1; * *P <* 0.05; ** *P <* 0.01; *** *P* ≤ 0.001.

An analysis of directional connectivity with GC revealed a confirmatory but much more fine-grained picture with KO-induced aberrations in all four frequency bands depending on the connection and direction (Fig. 2M-O). Most prominently, we found strongly elevated theta-range GC for all projections departing in either subdivision of the hippocampus. This confirms the hippocampal (as opposed to prefrontal) origin of the theta hyper-connectivity phenotype in *Gria1*-knockout mice, that we had postulated before based on the normalization of this phenotype in mice with hippocampal rescue of GluA1-expression [42]. Likewise, beta and gamma dHC➔PFC GC was strongly reduced in knockouts (Fig. 2M), in line with reduced phase-synchronization measures (Fig. 2D, J), while PFC➔dHC beta and gamma GC were even mildly elevated. This again suggests a hippocampal origin of the reduced coupling in this frequency range. The most prominent GC was found in the delta range, with PFC➔d/vHC GC being significantly larger than the delta GC in the opposite direction in both genotypes. However, there was no consistent alignment between genotype-related differences in the wPLI and in the GC measures in the delta-range.

In contrast to the overall relatively redundant picture between coherence, wPLI, and GC in the theta-range, the analysis of coupling using further activity parameters did not paint the same picture. Assessing SPC using the mean resultant length (MRL) of the vector representing average spike-occurrence in theta-phase space (Fig. 2P), we found that coupling of PFC and vHC spikes to dHC theta phase was not elevated (qualitatively even lower) in *Gria1*-knockouts. However, in line with the former measures, phase-locking of PFC and dHC spikes to vHC theta was increased in KO mice (Fig. 2Q).

Likewise, when analysing the gamma-theta cross-regional PAC, dHC-gamma was coupled stronger to vHC-theta – and also to PFC-theta - in knockouts (Fig. 2R) in line with the MRL metric. But there was also an increased coupling of vHC-gamma to the theta-oscillations in both other regions that had no correspondence to the SPC analysis. PFC-gamma to vHC-theta coupling was even marginally *reduced* in knockouts which contrasts sharply with the results from all other measures (Fig. 2R).

To complete our analysis of directed metrics, we investigated four measures of consistent phase-differences between potentially coupled regions. Firstly, we examined the imaginary component of the coherence metric (ImC). This showed a characteristic ~90°-shift between the theta, beta, and gamma oscillations of the PFC *vis-à-vis* the dHC, that was however equal between genotypes. Similar but smaller shifts were seen for theta in the other connections (Fig. 3A). Genotype-related differences were only seen in the PFC-vHC connection in the theta and, inversely, the gamma range (Fig. 3A). Furthermore, we investigated two directional measures obtainable from the SPC: the average theta-phase of the mean resultant vector (MRV; [36]) and a cross-correlational analysis of incremental shifts of the MUA relative to the theta-cycle [32]. The MRV of PFC and vHC spikes relative to the dHC theta phase were significantly shifted between genotypes: while they occurred during the rising phase of theta in knockouts, they occurred at its through or falling phase in wildtype mice (Fig. 3B). Similar shifts of PFC/vHC spikes relative to dHC theta were seen with the cross-correlational analysis, although a significant difference between genotypes was only detectable in the vHC(spikes)-dHC(theta) coupling (Fig. 3C). In the PFC-vHC connection, the spiking in one region always led the theta-oscillation of the other region, irrespective of direction or genotype (Fig. 3C).

**Figure 3.**
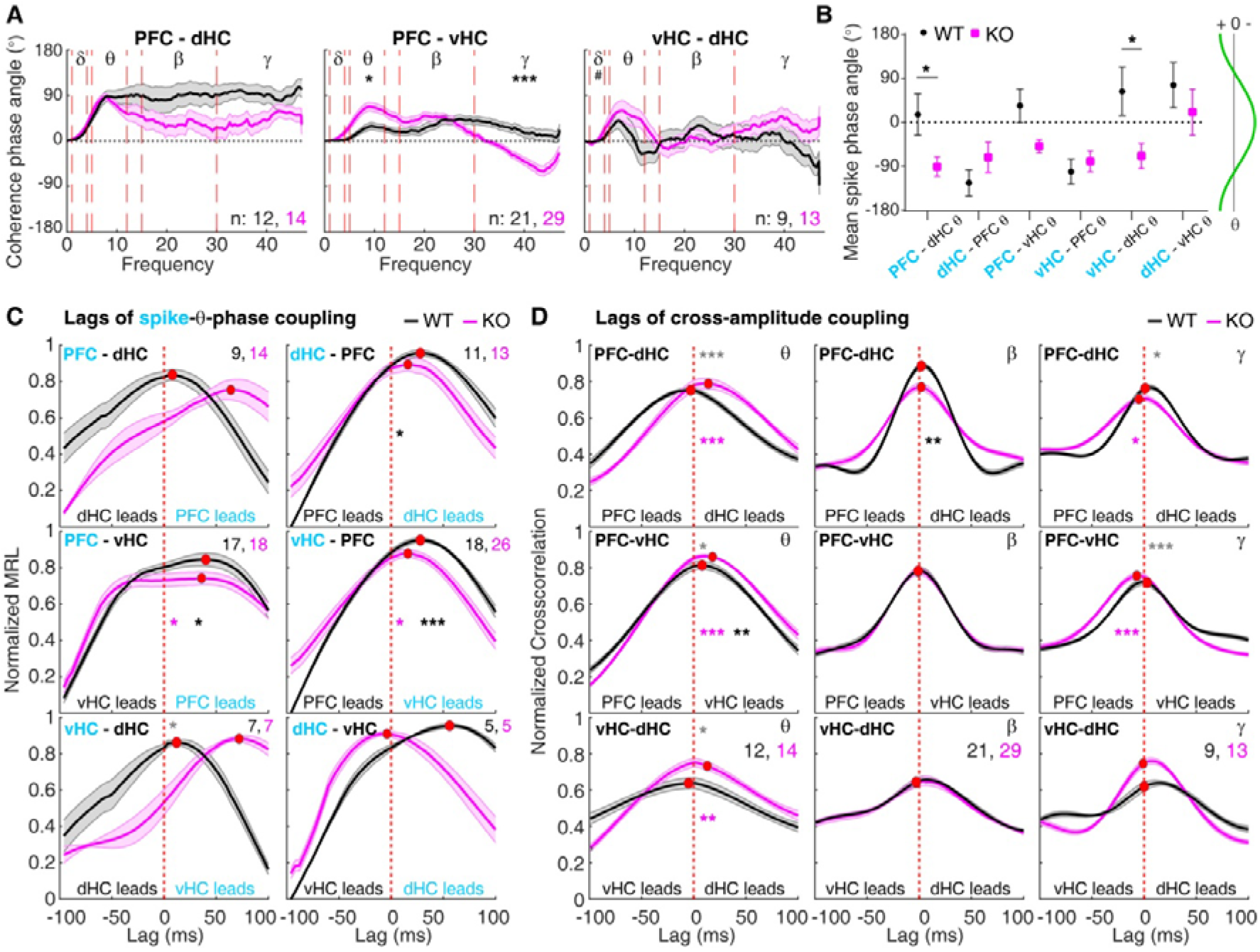
Multi-parameter assessment of phase-shifts and lags within inter-regional coupling in *Gria1*^−/−^ and wildtype controls across 10 min novelty-induced activity. (**A**) Spectra of imaginary part (phase angle) of coherence along the connection identified above each panel. Red lines in spectra indicate the boundaries of the analysed frequency bands named by the greek letters at the top. Stars indicate significant differences between genotypes (*t*-test). (**B**) Average theta-phase of the mean resultant vector resulting from the SPC analysis. Stars illustrate significant group difference (Watson-Williams test). The phase of theta corresponding to the degree-value are shown on the right (horizontal axis illustrates voltage of LFP). (**C**) Cross-correlation functions between spike-times and instantaneous amplitudes for the directional connections identified at the top of each panel (region contributing spikes in blue). Note that some data has been excluded based on lag-amplitudes above 100 ms; hence, contributing *N*-numbers for both genotypes (colour-coded) are shown in each panel. No genotype-related differences have been detected (*t*-test); black stars refer to a significant difference of the lag (temporal shift) from 0 ms in WT mice (Wilcoxon sign-rank test) and none of the shifts were significantly different from 0 in the KO-group. (**D**) Cross-correlation functions of instantaneous amplitude curves in the connections and frequency bands named at the top of each sub-panel with peak values indicated by a red dot. Statistical indictors in black (WT) or purple (KO) in (C) and (D) refer to a significant difference of the lag (temporal shift) from 0 ms (Wilcoxon’s signed rank test). Grey indicators refer to pair-wise comparisons between genotypes (*t*-test), which are, however, to be interpreted with caution, given that these method are designed to identify shifts away from 0, not to quantify the lag-amplitude. Solid lines display mean and shaded area SEM throughout. * *P <* 0.05; ** *P <* 0.01; *** *P* ≤ 0.001.

The analysis of significant non-zero lags by amplitude cross-correlation of LFP oscillations painted a rather different picture compared to the three analyses of phase-shifts before: here, genotype-related differences in theta-range directionality were found in all three connections. Also, vHC theta was leading PFC theta in both genotypes, while the dHC-theta lead theta oscillations in the other two regions only in the *Gria1*-knockouts (Fig. 3D). In reverse, in the gamma range, PFC led both hippocampal regions exclusively in the knockouts, which matches neither the expectation from the GC nor the ImC analysis.

In summary, while the identification of genotype-related differences in coupling is similar between some measures (especially coherence and GC), there is also a considerable lack of redundancy across the different measures of interregional connectivity (see overview in Table 1).

**Table 1.** Pairwise comparison between wildtype and *Gria1*-knockouts. Overview over genotype-related statistical comparisons of the data displayed in Figures 2-3 (average metric) and 4 (slope metric). GC results are derived from Sidak post-hoc test after repeated-measures ANOVA across both directions of a connection and genotypes; MRL-Phase is compared with the Watson-Williams test [36]; all other *P*-values are derived from independent-sample *t*-tests. For LFP-based measures (coherence, wPLI, GC) the *P*-values in the theta-range refer to peak-theta (not mean-theta). Arrows in directional measures indicated direction of coupling, → direction labelled in column name (e.g. PFC→dHC in the PFC-dHC column), ← opposite direction. For MI and MRL measures, the region named first corresponds to the region that contributes the theta-oscillation to the analysis. White background, no measure available or assessed; grey background alone, *P* ≥ 0.1; # *P* < 0.1; * *P* < 0.05, ** *P* < 0.01, *** *P* ≤ 0.001.

| KO vs. WT |  | delta |  |  | theta |  |  | beta |  |  | gamma |  |  |
| --- | --- | --- | --- | --- | --- | --- | --- | --- | --- | --- | --- | --- | --- |
|  |  | PFC-dHC | PFC-vHC | vHC-dHC | PFC-dHC | PFC-vHC | vHC-dHC | PFC-dHC | PFC-vHC | vHC-dHC | PFC-dHC | PFC-vHC | vHC-dHC |
| average metric | Coherence |  |  |  | *** | *** | *** |  |  |  | * |  |  |
|  | wPLI | ** |  | * | * | *** | ** | ** | # |  | * | * | # |
|  | GC → |  |  |  | # |  | * | # |  |  | * | # | * |
|  | GC ← |  | * | * | ** | *** | * | *** |  |  | *** |  | # |
|  | MRL → |  |  |  | # |  | * |  |  |  |  |  |  |
|  | MRL ← |  |  |  |  | ** |  |  |  |  |  |  |  |
|  | MI/PAC → |  |  |  | * | ** | * |  |  |  |  |  |  |
|  | MI/PAC ← |  |  |  | * | # | * |  |  |  |  |  |  |
|  | ImC |  |  | # |  | * |  |  |  |  |  | *** |  |
|  | CC |  |  |  | *** | * | * |  |  |  | * | *** |  |
|  | MRL-Phase → |  |  |  | * |  | * |  |  |  |  |  |  |
|  | MRL-Phase ← |  |  |  |  |  |  |  |  |  |  |  |  |
|  | MRL-Lag → |  |  |  |  |  |  |  |  |  |  |  |  |
|  | MRL-Lag ← |  |  |  |  |  | * |  |  |  |  |  |  |
| slope metric | Coherence | *** | ** | * | ** | *** | # | ** |  |  |  | # |  |
|  | wPLI | # |  |  |  | ** |  |  |  |  |  |  |  |
|  | GC → | ** | * |  | ** |  | * | * |  |  |  | ** |  |
|  | GC ← |  | ** | ** |  | *** | * |  |  |  |  |  | * |
|  | MI/PAC → |  |  |  |  | ** |  |  |  |  |  |  |  |
|  | MI/PAC ← |  |  |  |  | * | * |  |  |  |  |  |  |
|  | CC |  |  |  | * |  |  | # |  |  |  |  | * |

### Differences in detecting increases of inter-regional coupling over time in Gria1-knockouts across measures

As a second indicator for redundancy between connectivity measures, we investigated potential physiological correlates of the characteristic divergence of exploratory drive between the two genotypes over time (Fig. 1B-C). This divergence is likely induced by a failure of spatial short-term habituation in *Gria1*-knockout mice resulting in *increasing* exploration - as opposed to decreasing activity seen in controls [42, 43]. To allow for an efficient analysis, we captured the change of a given parameter over time in a single number, namely the slope of the linear interpolation across the time series over the 10 min of the test, as exemplified for three measures in Fig. 3A, C and E. We previously found that both local theta power in the dHC and also dHC-PFC theta coherence displayed a characteristic divergence between the groups, that mirrored exploratory behaviour [42]. In the current analysis, this pattern emerged much more broadly across multiple power and coherence measures, including in local PFC power in all analysed frequency bands, as well as in gamma and (at trend-level) theta-peak power in the hippocampal regions (Fig. 4A-B). For coherence, the KO-related increase in slopes was limited to the delta and theta range and was apparent in the hippocampal-prefrontal connections (confirming our earlier results) and marginally for intra-hippocampal coupling (Fig. 4C-D). In the beta and gamma range either no group-difference occurred or – for PFC-dHC beta coherence – it was even inversed with a higher slope in wildtype mice. Stunningly, this pattern was not reproduced by the wPLI analysis (Fig. 4E-F) – even in the one case where the coupling slope was increased in knockouts in *both* metrics (PFC-vHC, theta), the metrics differed in the respect that, in wildtype controls, theta-wPLI remained constant, while theta-coherence decreased over time.

**Figure 4.**
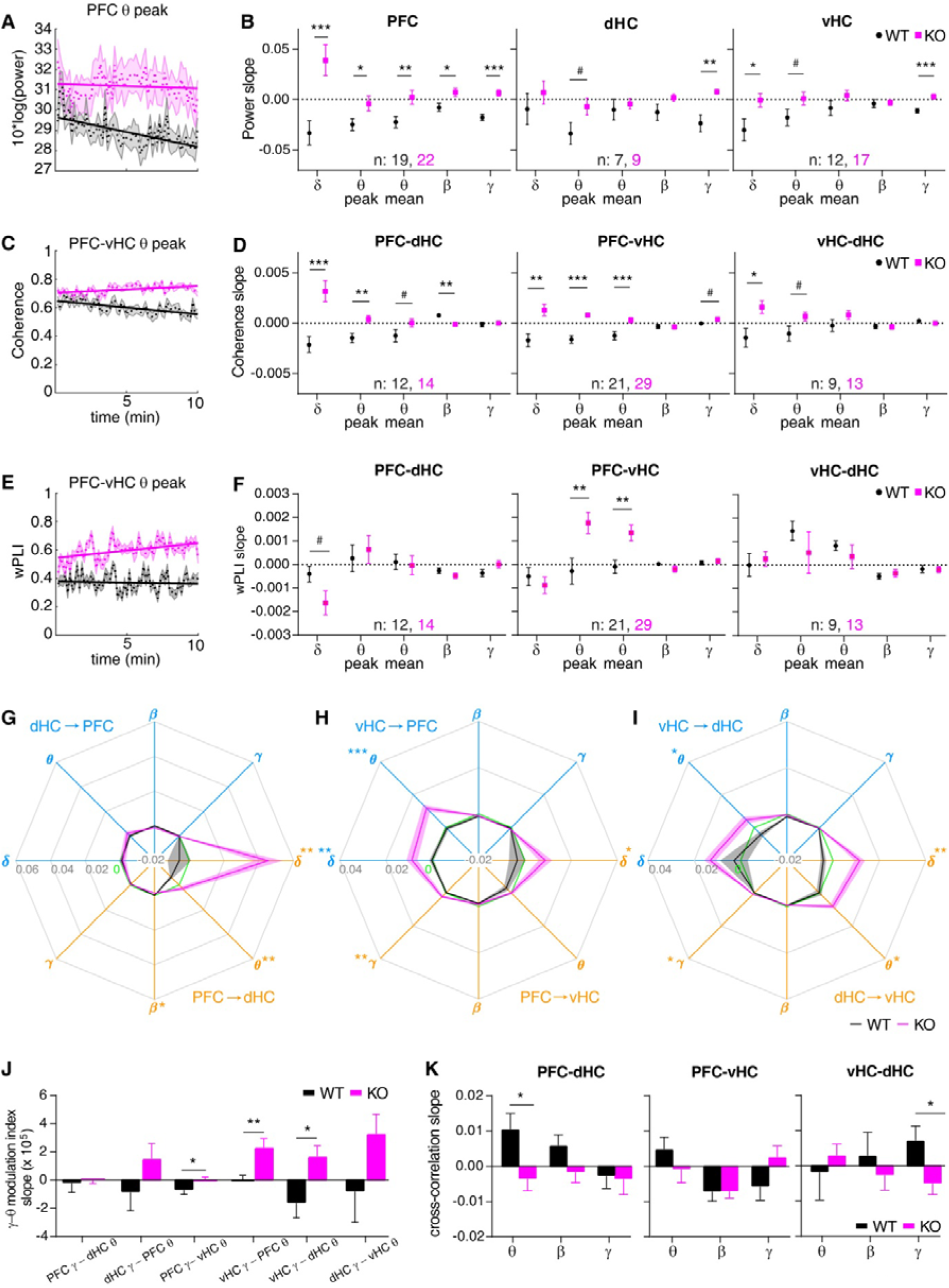
Changes of power and coupling strength over time during the 10 min open-field test. (**A, C, E**) Examples of individual measures of power (A), coherence (B), and wPLI (C) as they behave as population overage over the 10 min of novelty-induced activity in the open field (dashed line, mean; shaded area, SEM) with linear interpolation between time points overlaid (solid line) to determine the slope as indicator of temporal changes. (**B, D, F**) Average slope (temporal change) of power (B), coherence (D) and wPLI (F) in the indicated regions or connections (top of sub-panel) and frequency bands (x-axis). (**G, H, I**) Slope of GC in the frequency bands indicated by greek letters and along the directional connections identified by the colour (blue: dHC➔PFC (G), vHC➔PFC (H), vHC➔dHC (I); orange: reverse of the before). Statistical indicators in the same colour identify a difference between genotypes (*t*-test). (**K**) Slope of theta-gamma PAC in the stated directional connections. (**J**) Slope of cross-correlation lags indicating putative changes of temporal shifts of the oscillations in the named frequency bands. Black stars indicate significant differences between genotypes (*t*-test), and error bars or shaded regions indicate SEM throughout. # *P* < 0.1; * *P <* 0.05; ** *P <* 0.01; *** *P* ≤ 0.001.

GC remained largely constant or slightly decreased over time in wildtype mice, irrespective of connection or frequency band (Fig. 4G-I). In *Gria1*^−/−^ mice, in contrast, GC *increased* over time in the delta and theta range in most connections leading to genotype-related differences in the vHC➔PFC (δ θ), vHC➔dHC (θ), dHC➔vHC (δ θ), PFC➔vHC (δ γ), and PFC➔dHC (δ θ) projections. Thus, except for an isolated match in the vHC➔PFC theta-connectivity, the GC metric does not align with wPLI-based slope assessment, but provides a near perfect match to the coherence slope pattern (Table 1). The latter observation even extends to the one instance of PFC-dHC beta coupling where the slope is higher in wildtype than in KO mice (Fig. 4G-I). While the slope was not calculated for SPC because this measure requires a considerable number of spikes, the slope of the gamma-theta PAC could be determined. Here again, the expected divergence between an increasing slope in the PAC-strength in knockout contrasted with a constant or qualitatively decreasing slope in wildtypes, whereby the individual results partly matched the pattern seen with GC: a divergence at the intra-hippocampal projections bidirectionally and at the coupling of hippocampal (dHC/vHC) gamma to PFC theta (Fig. 4G-J). Cross-correlational lags, in contrast, did not change in any pattern that resembled the other measures (Fig. 4K).

### Lack of redundancy between coupling measures revealed by bivariate correlation analysis

Given that the above analysis of comparing genotype-related differences across measures ultimately allows only a *qualitative* judgement about the epistemiological redundancy of interregional measures, we supplemented our analysis by a more quantitative analysis in form of bivariate correlations between pairs of parameters and within genotypes and connections using the average value for each parameter in each electrode pair as dependent variable. This revealed multiple levels of complexity when analysing the relation between the metrics. On the one hand, at the level of isolated observations, the correlations supported the commonalities between measures already seen with the two prior analyses. For example, PFC-dHC theta coherence correlated strongly with dHC➔PFC theta-GC in wildtypes (Fig. 5A). However, this correlation did neither exist in the *knockouts* in the same connection (Fig. 5A) nor in the same genotype but the PFC-*vHC* connection (Fig. 5B). Indeed, PFC-vHC theta coherence, did correlate highly with GC in the opposite, i.e. PFC➔vHC, direction but not in the vHC➔PFC direction – and it did so across all frequency bands (Fig. 5B), which was not the case in the other two connections (Fig. 5A, 6A). In general, when carefully examining each pair of metrics, it becomes apparent that the correlation seen in one genotype and connection would rarely reappear in another one (Fig. 5A-B, 6A).

**Figure 5.**
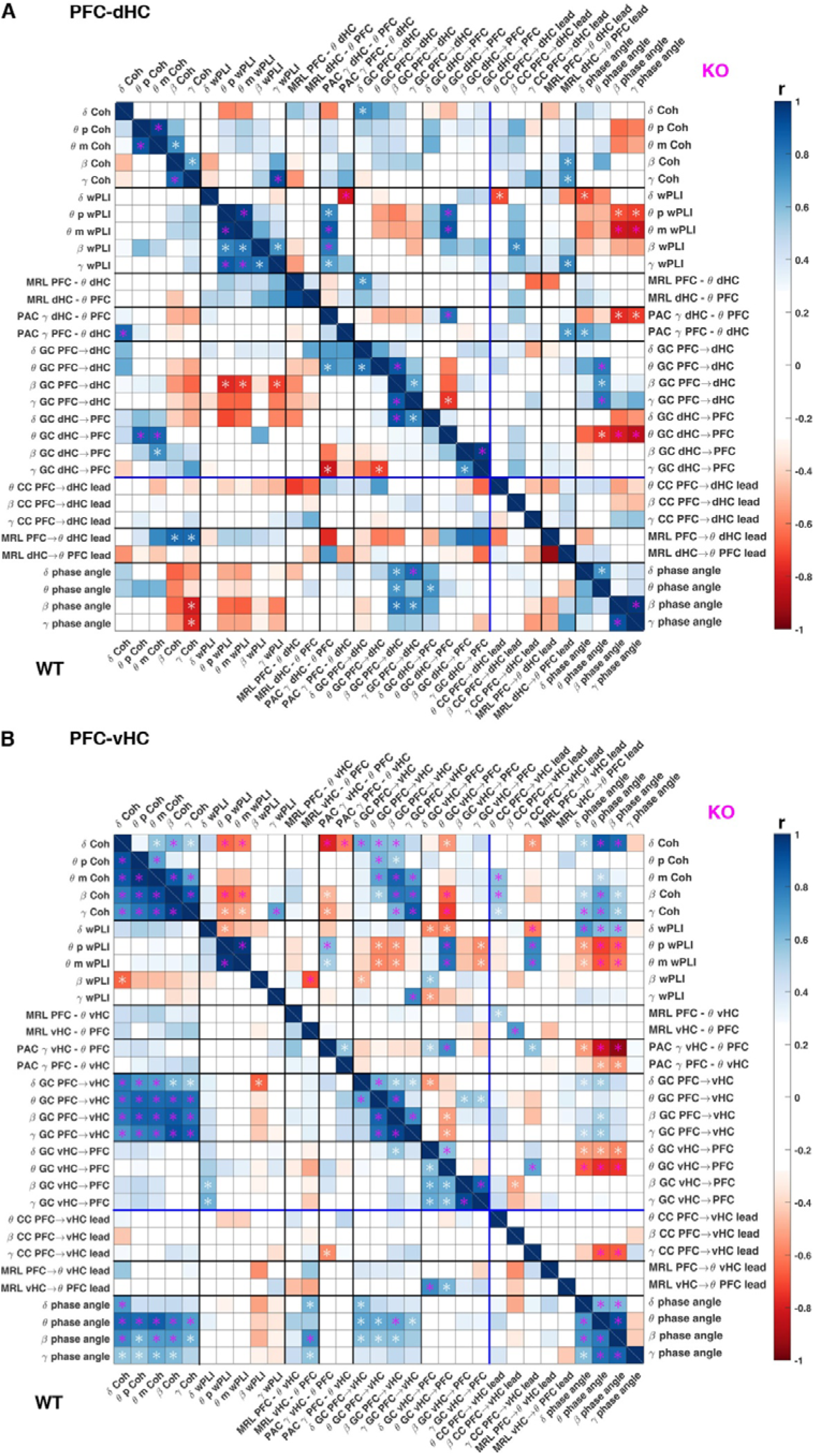
Correlations between individual measures of hippocampal-prefrontal connectivity. (**A, B**) Coefficient (r, colour of squares) and significance (star within squares) of bivariate correlations between individual measures of connectivity in the PFC-dHC (A) and PFC-vHC (B) connections within KO (top-right) and WT (bottom-left) mice. White stars, *P* < 0.01; purple stars, *P* < 0.001. Different classes of measures are separated by thicker lines. Measures of temporal shifts are separated from measures of connectivity by blue lines.

**Figure 6.**
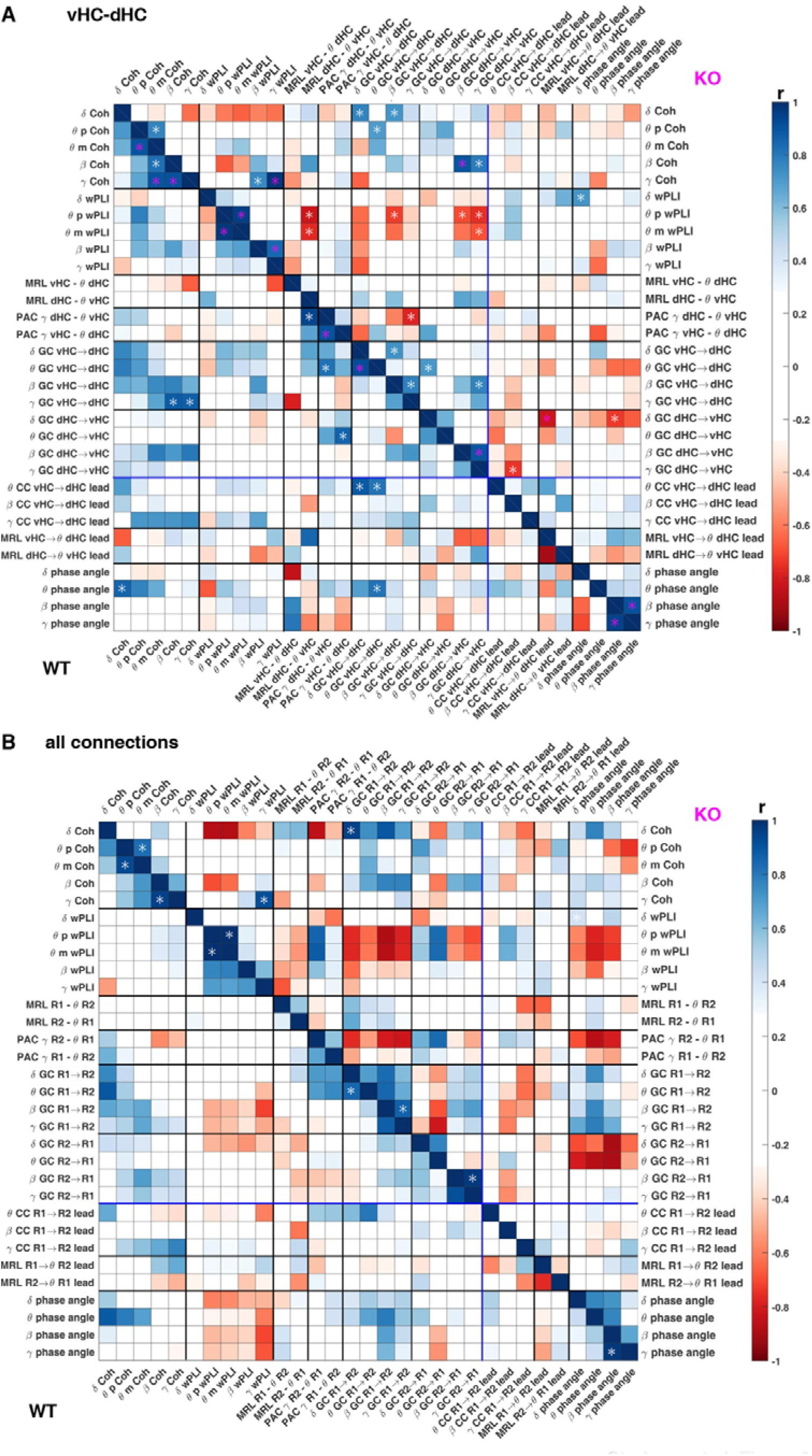
Correlations between individual measures of intra-hippocampal and overall connectivity. (**A**) Coefficient (r, colour of squares) and significance (star within squares) of bivariate correlations between individual measures of connectivity in the vHC-dHC connection within KO (top-right) and WT (bottom-left) mice. (**B**) Same display as in (A) but indicating the *average* correlation coefficient across the three connections (Fig. 5 A, B, Fig. 6A) by the colour of a square and significance only if a significant correlation existed in every one of the three connections. White stars, *P* < 0.01; purple stars, *P* < 0.001. Different classes of measures are separated by thicker lines. Measures of temporal shifts are separated from measures of connectivity by blue lines.

In order to evaluate this systematically, we calculated the average correlation coefficient for each pair across the three connections and indicated its significance only if it was given in all of them (Fig. 6B). In wildtype mice there was not a single pair of distinct metrics that achieved a significant correlation in all three connections-except for some expected correlations between neighbouring frequency bands within the same metric. The correlations that achieved a high average correlation coefficient (r > 0.7) are nevertheless notable and included correlations between the amplitude and the imaginary part (ImC) of delta and theta coherence, and also between theta GC and most other measures namely delta/theta coherence, theta ImC, gamma-theta PAC, and theta amplitude cross-correlations. SPC (MRL) and wPLI, in contrast, correlated with virtually no other measure. This synopsis largely aligns with the redundancy patterns seen with the two former analyses (Table 1).

However, it does largely not align with the results of the same analysis in the KO mice (Fig. 6B). Here, two consistently significant correlations were apparent – gamma wPLI to gamma coherence, and delta GC to delta coherence. When investigating the set of high average correlation values, a moderate correlation between GC and the amplitude and imaginary component of coherence as well as with PAC emerged again. In addition to these patterns-seen at least qualitatively also in wildtype mice - in knockouts also correlations between theta wPLI and three other measures emerged: gamma-theta PAC, theta GC, and beta amplitude cross-correlation (Fig. 6B). This illustrates, that even partial redundancies between measures seen in wildtype mice are still dependent on the genotype under investigation, and are hence not reflecting *a priori* redundancies.

### Sensitivity of measures to reference location

The choice of placement site for the reference electrode varies considerably between studies, and both the referencing to the ground screw above the cerebellum (as done for all above analyses) and to the anterior part of the frontal cortex are widely used. In order to investigate the effect of this difference, we recorded a separate reference signal from a frontal reference screw and used it to digitally re-reference all recorded data by subtracting this signal from the recorded LFP traces before re-calculating local power, coherence, wPLI, and GC.

Using repeated-measure ANOVAs with the within-subject factor of re-referencing and the between-subject factor of genotype, we found that the location of the reference has quite a substantial influence on the results. There were significant effects of re-referencing on delta power, coherence, and GC in all brain regions (except for the MD) and connections, while the effect on wPLI was comparatively minor (but note that delta-wPLI is generally very low and entirely different from delta-coherence and GC; Fig. 7E-M). In the theta-range, in contrast, re-referencing affected power only in the dHC, but strongly impacted coherence, wPLI, and GC alike along both hippocampal-prefrontal connections – not only in terms of significant effects of re-referencing, but also in terms of genotype-reference interactions, which indicate that the prior conclusions on theta-range connectivity are partly dependent on the position of the reference. In the GC measure, interactions were apparent in the d/vHC➔PFC direction but not in the reverse (Fig. 7K-L). Nevertheless, there were also significant effects of genotype in those connections and measures, suggesting that the fundamental observation of elevated hippocampal-prefrontal theta-connectivity in knockouts still holds, especially for the PFC-dHC connection and the GC measure in general (Fig.8E-F,H-I,K-L). Intra-hippocampal theta-connectivity was not much affected by the reference placement, irrespective of measure (Fig. 7G,J). In the higher frequency-ranges the effects were more mixed. Beta power in the dHC and coherence – but only partly wPLI and GC – along its connections were affected by reference placement. In the gamma range, re-referencing impacted power in the PFC and dHC, wPLI in the PFC-d/vHC connections, and coherence along all three connections (Fig. 7A-J). In fact, the formerly observed lower PFC-dHC gamma-coherence and wPLI in knockouts (Fig. 2D, J) was dependent on the reference placement for detection (interaction effect only for coherence and wPLI, Fig. 7E,H). A similar observation holds for the PFC-vHC gamma connectivity which was increased in KOs in the wPLI, but not the coherence measure (Fig. 2E, K). Here again, an interaction indicated that the absence or presence of this difference in the coherence measure depends on the reference location (Fig. 7F), while an effect of genotype is maintained when using wPLI even though an interaction is found in addition (Fig. 7I). The impact of referencing on gamma-GC, in contrast, was limited to the dHC➔PFC projection (Fig. 7K-M).

**Figure 7.**
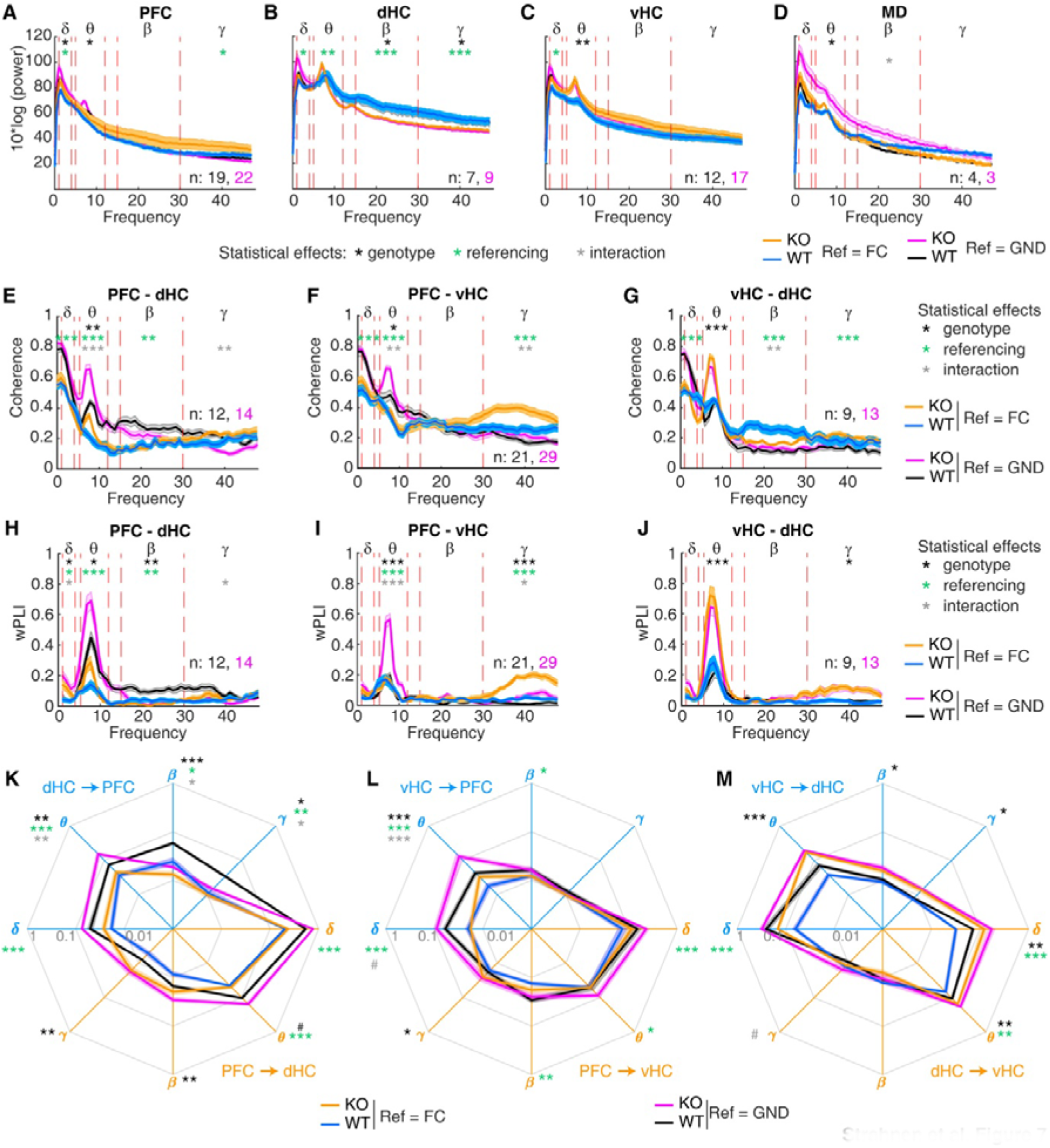
Assessment of the impact of the reference electrode placement on the measurement of power and connectivity. (**A-J**) Spectra for power (A-D), coherence (E-G) and wPLI (H-J) for the regions or connections indicated at the top of each panel, shown for standard referencing to the ground screw above the cerebellum (black, WT and purple, KO; as in Fig. 2) or digitally re-referencing to the reference screw above the frontal cortex (blue, WT and orange, KO). Red lines indicate the boundaries of the analysed frequency bands named by the greek letters at the top. (**K-M**) GC in the frequency bands indicated by greek letters and along the directional connections identified by the colour (blue: dHC➔PFC (K), vHC➔PFC (L), vHC➔dHC (M); orange: reverse of the before; display as in Fig. 2M-O). Throughout, shaded regions indicate SEM and stars indicate results of RM-ANOVA: black, effect of genotype; green, effect of chosen reference; grey, genotype-reference interaction. In the theta-range the statistics for coherence and wPLI refer to peak-theta. * *P <* 0.05; ** *P <* 0.01; *** *P* ≤ 0.001.

In summary, a frontal reference screw - as often used when studying prefrontal-hippocampal connectivity [37, 38] - may considerably alter the results obtained for LFP-based measurements of connectivity between the PFC and the hippocampus. Somewhat surprisingly, the wPLI measure does not eliminate this contingency but only reduces it, especially in the beta-gamma range. Effects on GC are particularly visible in the low (delta/theta) frequency range and (as interactions) in the direction from hippocampus towards PFC.

## Discussion

We here examined the level of redundancy and experimental contingencies of the most widely applied measures of interregional directed and non-directed neuronal connectivity that are obtainable when recording LFP and MUA with chronically implanted field electrodes in awake rodents. This analysis revealed a surprisingly large absence of expected redundancies between such measures and a worrying contingency with respect to the location of the reference electrode. This suggests that the implicitly held belief that experimental results obtained with one metric of connectivity and one configuration for referencing would allow *general* conclusions about aberrations in inter-regional functional connectivity is problematic. Intriguingly, a similar conclusion has been reached by a recent study on connectivity measures applied on human EEG data [44]. Even though the usage of single-unit activity from electrode bundles or arrays [37, 38] will certainly improve the assessment of SPC and its related directional measures beyond what was achievable in our setting with MUA, this will unlikely resolve this fundamental problem regarding redundancy. A particular analytical problem for this area of study appears to be the lack of benchmarking of the sensitivity, specificity, and robustness of the individual measures against a ground truth of actual physiological trans-synaptic activity along anatomically verified connections. Instead, the applied metrics have been largely introduced based on either theoretical considerations or discoveries of biological phenomena of coupling, and may be maintained due to historical path dependency. In absence of such benchmarking and while facing considerable logistical limits in applying multiple referencing and multiple metrics for every experiment, our analysis at least qualitatively implies some guidelines on how to maximize the analytical gain from similar experiments.

Firstly, our correlation analysis implies that parametric Granger Causality would be the measure of choice when investigating inter-regional connectivity purely based on LFP signals. This is because GC-values correlated with the majority of other measures to some extent, especially in wildtype mice (Fig. 5, 6). Likewise, genotype-related differences seen in the other connectivity measures (Fig. 2-4) were largely mirrored by aberrations in GC. Finally, as a directed measure, GC renders a more fine-grained picture of connectivity than other measures. For these three reasons, GC seems to be the metric with the lowest risk that true underlying aberrations of connectivity go undetected. Secondly, when adding further metrics to the analysis, SPC and wPLI would be most complementary because they correlated least with the other measures, including GC, both in the bivariate correlation analysis within subjects and with respect to detecting genotype-related effects. Thirdly, for LFP-based measures, the reference electrode should be placed in a brain structure that is largely separate from the brain regions between which connectivity is studied. A frontal screw may easily obscure phenotypes in prefrontal connectivity as it may pick-up field potential signals from the PFC [5].

In summary, our analysis calls for a more cautious interpretation of previous findings in the rodent literature on inter-regional coupling (especially when regarding negative results), the need for better benchmarking of individual measures, and the necessity to report multiple measures of connectivity in future studies.

## Materials and Methods

### Animals

Male and female *Gria1* knockout (*Gria1^−/−^*, *Gria1*^tm1Rsp^; MGI:2178057) [45] mice (N = 20, 12 males) and wild-type littermate controls (N = 12, 9 males) were bred from heterozygous parents. Animals were group-housed in Type II-Long individually ventilated cages (Greenline, Tecniplast, G), enriched with sawdust, sizzle-nest^TM^, and cardboard houses (Datesand, UK), and subjected to a 13◻h light / 11◻h dark cycle. The mice were implanted with electrodes at ca. 9 mo of age and were tested in the open field test ca. 3-5 weeks later to allow recovery from surgery intermittently. All experiments were performed in accordance to the German Animal Rights Law (Tierschutzgesetz) 2013 and were approved by the Federal Ethical Review Committee (Regierungsprädsidium Tübingen) of Baden-Württemberg, Germany (licence number TV1399).

### Surgery

Electrode implantation surgeries under general isoflurane-anaesthesia and a broad peri-operative analgesic regime were conducted similarly as previously described [42]. Briefly, single polyimide-insulated tungsten wires of 50 µm diameter (WireTronic Inc., CA, US) were implanted, with reference to Bregma (in mm), into the PFC (AP +1.8-1.9, ML 0.3-0.35; 1.8-1.9 below pia), MD (AP −1.2, ML 0.3, 2.7 below pia), dHC (AP −1.9-2.0, ML 1.5, 1.4 below pia), and vHC (AP −3.1-3.2, ML 2.9-3.0, 3.4 mm for single and 3.8-3.9 mm for dual electrodes below pia). In a majority of mice, dual electrodes were used for PFC and vHC, whereby the second electrode was placed about 0.5 mm higher than the stated distance from pia. In later analysis, the data from each electrode was regarded as unit of observation (*N*), so that a single mouse could contribute up to an *N* = 4 for vHC-PFC connections, and up to an *N* = 2 for dHC-vHC, PFC-vHC, MD-PFC, and MD-vHC connections. Both hemispheres were implanted at roughly equal proportion. Stainless steel screws (1.2 mm diameter, Precision Technologies, UK) were implanted in the contralateral hemisphere ca. 1 mm from the midline above the cerebellum (AP −5.5) for ground and above the anterior frontal cortex (AP +4.0) for additional reference, and where connected with a 120 µm PTFE-insulated stainless steel wire (Advent Research Materials Ltd., UK; Fig. 1A). All electrode wires were connected to pins in a dual-row 6-pin or 8-pin connector (Mill-Max, UK) To later determine electrode placements *post-mortem*, electrolytic lesions were made after breathing ceased under terminal ketamine/medetomidine anaesthesia. Immediately afterwards, animals were transcardially perfused with PBS followed by 4% paraformaldehyde (PFA)/PBS and brains were post-fixed for 24 h in PFA/PBS. Coronal sections of 60 µm were cut on a vibratome in PBS and then washed 3 times in PBS, stained with DAPI, and mounted for inspection of lesion sites on an epifluorescence microscope (DM6, Leica).

#### Novelty-induced locomotion and recording

Animals were tethered to enable electrophysiology recordings and then placed into a novel environment consisting of a clear Type-III plastic cage (length 43 cm, width 22 cm, height 20 cm; Tecniplast) containing clean sawdust. Animals were allowed to explore for 10 min. The animals’ location in the open field was video-tracked with ANY-maze (Stoelting, UK) and the distance travelled was calculated in 2 min time bins. Prior to testing, a 32-channel RHD2132 headstage (Intan Technologies, CA, US) was plugged into the implanted connector via a custom-built adaptor that interfaced a 36-pin Omnetics connector (A79022-001, MSA components, G) with another 6-pin or 8-pin Mill-Max connector. The headstage was wired to an *Open-Ephys* acquisition board (https://open-ephys.org, US; obtained through the Open-EPhys store at Champalimaud, Portugal) via two light-weight flexible SPI-cables (Intan Technologies), daisy-chained through a custom-connected miniature slip-ring (Adafruit, NY, US). The adaptor was wired so that all signals were referenced to the ground-signal obtained from above the contralateral cerebellum, while the signal from the additional frontal reference screw was recorded separately (for later offline re-referencing) like the LFP channels, i.e. also referenced to ground. Using the RHD2132 headstage, the Open-Ephys acquisition board, and the Open-EPhys acquisition software, data were amplified and digitized, sampled at 10 kHz, and digitally high-pass filtered at 0.1 Hz for acquisition of raw-data (for MUA and GC analysis) and simultaneously band-pass filtered at 0.1 – 250 Hz (for all remaining analysis of LFP signals).

### Data processing and analysis

All signal analyses were done in MatLab (MathWorks). Data were exported to Matlab and, for all LFP analyses, down-sampled to 1 kHz and analyzed with custom-written scripts. To reduce low frequency drift, signals were first detrended using the *locdetrend* function of the Chronux signal processing toolbox (http://chronux.org/) with 1 s of data and a sliding window of 0.5 s.

### Spectral Analysis

Power and coherence spectra as well as the phase angles were calculated with Chronux routines implemented in the Chronux toolbox using the multitaper method [46]. Power values were expressed as 10*log_10_ values for all analyses and the range of frequencies was set from 0.1 to 48 Hz. A bandwidth of 0.2 Hz and a total of 220 tapers were used to calculate power and coherence over the course of the 10 min exploration time. To analyse the temporal development, power and coherence were also calculated in 10 s bins using a bandwidth of 1 Hz and 19 tapers.

### Weighted Phase Lag Index (wPLI)

To address the issue of volume conduction we calculated the weighted phase lag index (wPLI) [9] using the routines implemented in the FieldTrip toolbox [47]. The 10 min exploration time was divided into non-overlapping 1 s bins and padded to the next power of two. The complex cross-spectrum was computed using a Hann taper with a spectral smoothing of 0.5 Hz. For temporal analysis, wPLI was averaged for each minute of the 10 min period using the same spectral parameters.

### Phase-amplitude coupling

Cross-frequency coupling (CFC, [23]) was assessed using the measure of phase-amplitude coupling (PAC), the statistical relationship between the phase of a low-frequency and the amplitude of a high-frequency component, in a cross-regional analysis [6, 28]. The 10 min recording was split into 1 min bins during which the PAC was calculated using the Modulation Index (MI, [28, 48]. Briefly, time series data is first band-pass filtered in the desired frequency range, followed by a Hilbert transform using the MatLab function *hilbert* which calculates the real and imaginary part of the signal to obtain the instantaneous amplitude and phase at any given time point. Theta phases were binned into eighteen 20 s intervals and the mean gamma amplitude was calculated in each phase bin. The distribution across bins was assessed using the Kullback-Leibler divergence [49] and normalized between 0 and 1. The MI is close to zero if the mean gamma amplitude is uniformly distributed over the theta phases and close to one if the mean gamma amplitude is exceptionally higher within one phase bin [28].

### Cross-correlation of instantaneous LFP amplitudes

To determine whether one signal was leading or lagging the other, amplitude cross-correlations of instantaneous amplitudes of LFP oscillations between all brain regions were performed [14]. The 10 min period was divided into 10 s bins with a 95 % overlap. First, the two signals were band-pass filtered in the respective frequency range; the Hilbert transform was computed using the MatLab function *hilbert* to calculate the instantaneous amplitude and the envelope of the signal. The mean amplitude was subtracted, and the cross-correlation between the amplitudes of the two signals was calculated with the MatLab function *xcorr* over lags ranging from −100 ms to + 100 ms; the lag at which cross-correlation peaked was determined [14]. While lags below −100 ms or above 100 ms would have led to exclusion of the respective data point [50], no instances of such lags were found in our dataset. To determine if the obtained lags or leads significantly differed from zero, Wilcoxon’s signed rank tests were performed.

### Granger Causality

Granger Causality (GC) was calculated using the MVGC-toolbox [51]. GC mainly applies to stationary signals which means that the variances are not excessively changing over time [20, 52]. Therefore, the 10 min period was divided into 1 min bins and the in-built trial averaging function was used to calculate GC in non-overlapping 10 s sections to ensure reasonable stationarity [53–55]. The 1 min bins were used for the analysis of GC over time and then averaged to obtain a GC value for the whole 10 min testing period. Raw LFP data was sampled down to 250 Hz to ensure a reasonable model order for autoregressive modelling [21, 51, 56]. The model order was obtained using the Bayesian Information Criterion (BIC, [57] as it was shown to provide the best fit to electrophysiological data [51]. The model order was fixed across all animals and trials to obtain comparable results [58]. Non-prefiltered data were used because empirical analyses have shown that filtering time series data increases the VAR model order and leads to high variances making it unsuitable for GC analysis [56]. To obtain GC values for specific frequency bands we first computed GC up to the Nyquist-frequency and then integrated over the desired frequency range [56]. A permutation procedure implemented in the MVGC-toolbox was performed to test the null hypothesis that values obtained by GC estimation occurred by chance [20, 51].

### Spike-Phase Coupling

Multi-unit activity was extracted by high pass filtering the raw signal above 800 Hz and applying a threshold at 3.5 standard deviations from the mean. Spikes were excluded, if the threshold exceeding was longer than 2 ms, and if spikes occurred within 1 ms form each other. LFP of the second brain region was filtered between 5 and 12 Hz using the *eegfilt* – function of the EEGLAB-toolbox [59]. To account for speed-dependent waveform asymmetry in the theta oscillation, the theta phase was defined by linear interpolation between consecutive troughs within each cycle [60, 61]. Only periods in which the theta amplitude was above 0.25 standard deviations of its mean were included to ensure sufficient theta oscillations and prevent spurious phase-determination by the subsequent Hilbert transform. The number of spikes was fixed to 1000 for each recording to prevent spuriously high MRL values and fluctuations in the firing rate. Each spike was assigned a theta phase and the mean resultant vector length (MRL) was calculated as an indicator for the strength of coupling using the CircStat-Toolbox [37, 62]. The MRL gets close to one when the spikes are concentrated around a certain phase of the theta oscillation and approaches zero when they are uniformly distributed. Additionally, the phase angles of the mean resultant vector were used to quantify differences in phase-angles between genotypes, which were statistically assessed with the Watson-Williams test for two samples [36, 62].

To determine the directionality between multiunit activity and theta oscillations, phase locking was calculated for 50 different temporal offsets ranging from −100 ms to +100 ms in steps of 4 ms. If the MRL peaked at a positive offset, spikes were most strongly locked to the next theta cycle, suggesting that spiking activity drives theta [32]. A Wilcoxon’s signed rank test was applied to determine if the lag or lead was significantly different from zero.

### Statistics

Genotype-related differences within the same metric and frequency range were assessed by independent-sample *t*-test or, in the case of GC (Fig. 2), by Sidak paired post-hoc tests conducted after a significant effect of genotype or interaction in the prior repeated-measures (RM) ANOVA. Variability in the data is displayed as standard error of mean (SEM) throughout.

## Data availability

All raw data and analysis code are available from the corresponding authors upon reasonable request.

## Acknowledments

We are grateful to Drs. Torfi Sigurdsson (Goethe-University Frankfurt) and Lionel Barnett (University of Sussex) for advice on analysis procedures, to Dr. Rolf Sprengel (Max-Planck Institute for Medical Research, Heidelberg) for the provision of *Gria1*^−/−^ mice, and to Stefanie Schulz (Ulm University) for assistance with histology. This work was funded by the Else-Kroener-Fresenius/German-Scholars-Organization Programme for excellent medical scientists from abroad (GSO/EKFS 12; to D.K.), the Juniorprofessorship programme of Baden-Württemberg (to D.K.), the DFG (KA 4594/2-1; to D.K.), the Brain and Behaviour Research Foundation (BBR/NARSAD, to D.K.) and the Institute of Applied Physiology of Ulm University.

## Author contributions

D.S. and D.K. designed the study and wrote the manuscript which was revised by all authors. D.S. and S.K.T.K. performed experiments. D.S. performed the data analysis, supported by A.M.B.

